# Effect of interglomerular inhibitory networks on olfactory bulb odor representations

**DOI:** 10.1101/2020.03.03.975201

**Authors:** Daniel Zavitz, Isaac A. Youngstrom, Alla Borisyuk, Matt Wachowiak

## Abstract

Lateral inhibition is a fundamental feature of circuits that process sensory information. In the mammalian olfactory system, inhibitory interneurons called short axon cells comprise the first network mediating lateral inhibition between glomeruli, the functional units of early olfactory coding and processing. The connectivity of this network and its impact on odor representations is not well understood. To explore this question, we constructed a computational model of the interglomerular inhibitory network using detailed characterizations of short axon cell morphologies taken from mouse olfactory bulb. We then examined how this network transformed glomerular patterns of odorant-evoked sensory input (taken from previously-published datasets) as a function of the selectivity of interglomerular inhibition. We examined three connectivity schemes: selective (each glomerulus connects to few others with heterogeneous strength), nonselective (glomeruli connect to most others with heterogenous strength) or global (glomeruli connect to all others with equal strength). We found that both selective and nonselective interglomerular networks could mediate heterogeneous patterns of inhibition across glomeruli when driven by realistic sensory input patterns, but that global inhibitory networks were unable to produce input-output transformations that matched experimental data and were poor mediators of intensity-dependent gain control. We further found that networks whose interglomerular connectivity was tuned by sensory input profile decorrelated odor representations more effectively. These results suggest that, despite their multiglomerular innervation patterns, short axon cells are capable of mediating odorant-specific patterns of inhibition between glomeruli that could, theoretically, be tuned by experience or evolution to optimize discrimination of particular odorants.

**Significance Statement:** Lateral inhibition is a key feature of circuitry in many sensory systems including vision, audition, and olfaction. We investigate how lateral inhibitory networks mediated by short axon cells in the mouse olfactory bulb might shape odor representations as a function of their interglomerular connectivity. Using a computational model of interglomerular connectivity derived from experimental data, we find that short axon cell networks, despite their broad innervation patterns, can mediate heterogeneous patterns of inhibition across glomeruli, and that the canonical model of global inhibition does not generate experimentally observed responses to stimuli. In addition, inhibitory connections tuned by input statistics yield enhanced decorrelation of similar input patterns. These results elucidate how the organization of inhibition between neural elements may affect computations.

## Introduction

Lateral inhibition is a fundamental feature of circuits that process sensory information (Isaacson and Scanziani, 2011). Computations mediated by lateral inhibitory circuits are diverse and can vary depending on the connectivity between neural circuit elements (Hartline et al., 1956; Yoshimura and Callaway, 2005; Poo and Isaacson, 2009; Kim et al., 2018; Znamenskiy et al., 2018). Lateral inhibition also plays a key role in the initial processing of olfactory information. In mammals, the first stage of this processing occurs among glomeruli of the olfactory bulb (OB), where olfactory sensory neurons (OSNs) expressing the same odorant receptor converge onto mitral/tufted cells - the principal OB projection neuron - as well as several classes of inhibitory interneurons (Mombaerts et al., 1996; Wachowiak and Shipley, 2006). Inhibition between glomeruli is a canonical feature of the OB network, with two major circuits mediating interglomerular inhibitory interactions. One circuit involves reciprocal dendrodendritic interactions between mitral/tufted and granule cells (Wachowiak and Shipley, 2006). Another circuit involves a class of inhibitory interneuron - short axon cells (SACs) - that branches extensively in the glomerular layer and regulates excitability of mitral/tufted cells in response to sensory input (Aungst et al., 2003; Whitesell et al. 2013; Banerjee et al., 2015; Liu et al., 2016). SACs thus constitute the first stage of lateral inhibition in the olfactory system.

Several recent studies have suggested that the SAC network mediates a form of global center-surround lateral inhibition that results in gain control that scales with the intensity of sensory input, due to the extensive branching of SACs, which can extend for up to 1 mm across the glomerular layer (Aungst et al., 2003; Cleland and Sethupathy, 2006; Banerjee et al., 2015). However, other studies have found that inhibition between glomeruli is more selective, with suppression of mitral/tufted cell outputs occurring in an odorant- and glomerulus-specific manner (Yokoi et al., 1995; Fantana et al., 2008; Economo et al., 2016). Similar examples of selective interglomerular inhibition have been explored in the olfactory system of nonmammalian species, where it is thought to enhance the discriminability of similar odorants (Linster et al., 2005; Wiechert et al., 2010; Mohamed et al., 2019). How – or whether - selective interglomerular inhibition arises from a widespread SAC network in the mammalian OB remains unclear.

SACs in fact constitute a morphologically diverse population that shows varying degrees of branching within a subset of glomeruli (Kiyokage et al., 2010). Depending on the nature of this connectivity, the SAC network could mediate selective interglomerular inhibition and odorant response sharpening, global inhibition and intensity-dependent gain control, or an alternate transformation of sensory input patterns. We approached this question computationally, by asking whether an interglomerular network derived from the known branching patterns of SACs is capable of mediating selective lateral inhibition of glomerular output and how the nature of SAC interglomerular connectivity impacts how this circuit transforms odorant representations. We constructed a simplified interglomerular network model based on quantitative, anatomical reconstructions of SAC glomerular innervations (Kiyokage et al., 2010) and in which a single parameter governed the selectivity of interglomerular inhibitory connections. Using experimentally measured OSN responses to odorant as inputs to the network and simulated mitral/tufted cell outputs from each glomerulus, we explored how odorant representations are affected by SAC network selectivity.

We found that interglomerular networks based on SAC morphologies are in fact capable of producing heterogeneous, glomerulus-specific inhibition when driven by realistic OSN input patterns. We also found - surprisingly – that globally connected networks were unable to produce input-output transformations that matched experimental data and were poor mediators of intensity-dependent gain control. Finally, we found that SAC networks with connectivity tuned by sensory input profile decorrelated odorant representations more effectively than randomly connected networks. These results suggest that, despite their multiglomerular innervation, SACs are capable of mediating heterogeneous patterns of inhibition between glomeruli that could, in theory, be tuned to optimize discrimination of particular odorants.

## Methods

### Network Connectivity

The models used in this study represent networks of nodes (glomeruli) connected by short axon cell (SAC)-mediated inhibitory connections. The connections of individual SACs across the network were based on previously-reported anatomical characterizations of SAC morphologies (Kiyokage et al., 2010). Interglomerular connectivity (connections between nodes in the network) resulted from the connections made by its constituent SACs. For the two network instantiations (containing 105 or 94 glomeruli, respectively), each glomerulus contained 40 SACs randomly sorted into two groups - oligoglomerular and polyglomerular - with probabilities of 0.8 and 0.2 respectively (Kiyokage et al., 2010). Each oligoglomerular SAC connected to 4 glomeruli and each polyglomerular SAC connected to 20. We modeled the selectivity of SAC networks by assigning each glomerulus a target set of glomeruli of size *m* (Fig. 1A) – a random set of glomeruli that its SACs were allowed to innervate. Thus, the magnitude of *m* determined the selectivity of interglomerular connections in the network, and the selectivity could be varied parametrically. We modelled the nonuniform distribution of innervation degree of SACs across their target set by giving each connection made by a SAC a random weight drawn from an exponential distribution with mean β=1.25. The total strength of connection from glomerulus *i* to glomerulus *j* (node *i* to node *j* in the network model), *w*_*ij*_, was the sum of the weights of the connections made by the SACs in *i* and innervating *j*. Networks with global, uniform connectivity were constructed by connecting each glomerulus to all other glomeruli with the same mean strength of interglomerular connections as described above.

**Figure 1.**
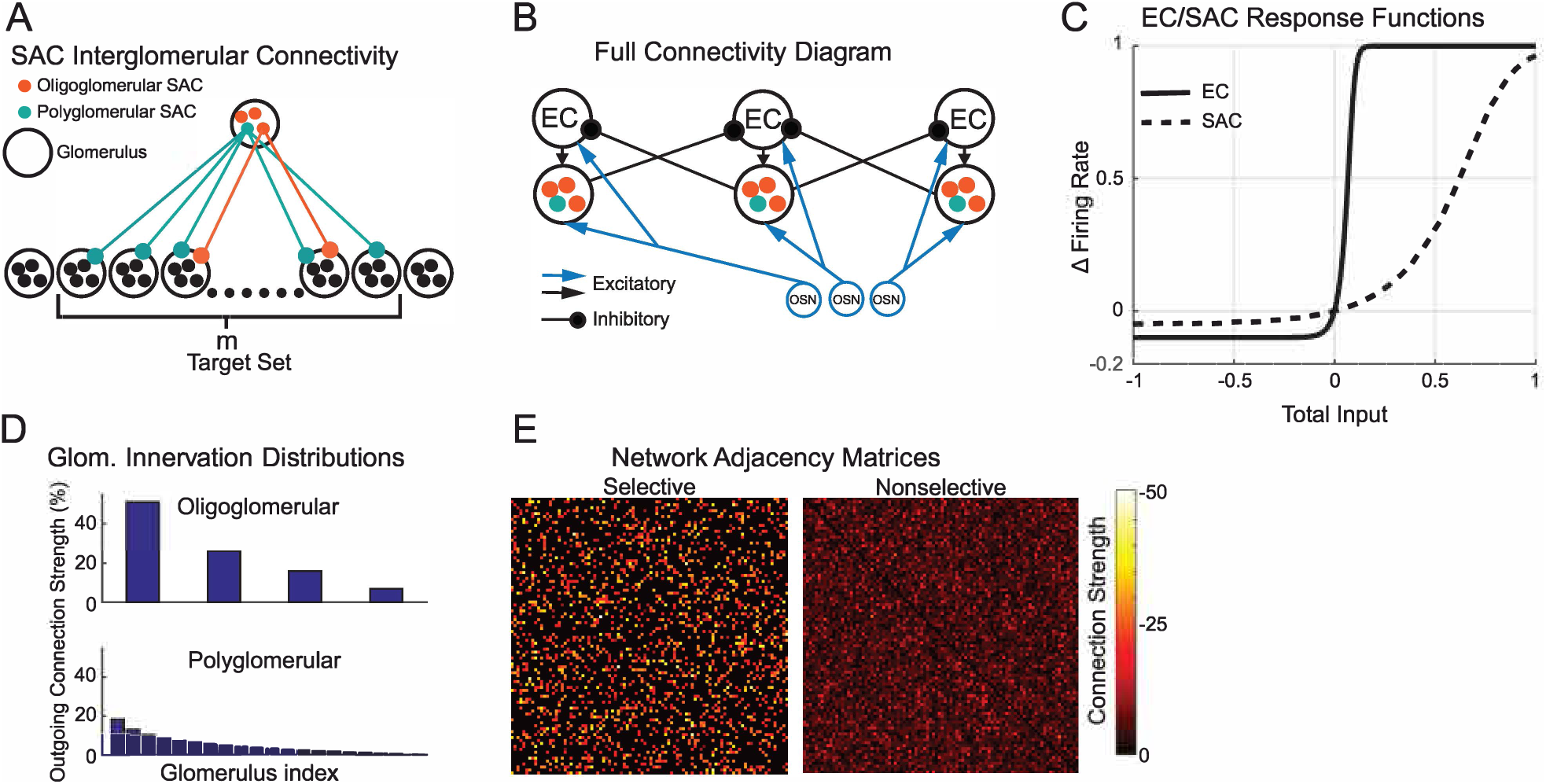
SACs Mediate Interglomerular Connectivity in a Simplified Olfactory Bulb Model. **A**. Diagram showing interglomerular connectivity. Oligoglomerular (orange) and polyglomerular (teal) SACs (short axon cells) form directed connections from their parent glomerulus to glomeruli in its target set. Oligoglomerular SACs form few connections compared to polyglomerular SACs but are much more common. **B**. The full connectivity diagram of SACs, ECs (excitatory cells) and OSNs. Excitatory connections are represented by arrows, inhibitory connections are represented by circles. **C** The responses of ECs and SACs to total input (excitation and inhibition) as measured by change in firing rate from baseline. **D**. The percentage of SACs’ total connection strength each innervated glomerulus receives. **E**. Left: the adjacency matrix of a selective (*m*<<*n*) network with few, strong inhibitory connections between glomeruli. Right: the adjacency matrix of an nonselective (*m*≈*n*) network with many weak inhibitory connections.

### Network Activity

After each network was constructed, responses to olfactory sensory neuron (OSN) inputs were computed using a simplified wiring scheme where a single excitatory cell (EC) was associated with each glomerulus, as was a single representative inhibitory SAC. Both cell types received excitatory input from the OSN associated with that glomerulus. Each node’s EC excited the SAC in its glomerulus and the SAC inhibited the ECs in other glomeruli using the network structure described above (Fig. 1B). Each EC and SAC had an activity level ranging between −0.1 to 1 and −0.05 to 1 respectively, where zero corresponded to the spontaneous firing rate, 1 corresponded to its maximal firing rate, and negative values corresponded to firing at rates below spontaneous (i.e., suppression). Changes in activity of the EC and SAC were governed by the sigmoidal functions (Fig. 1C):

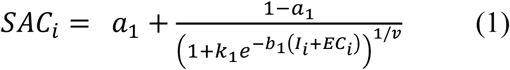

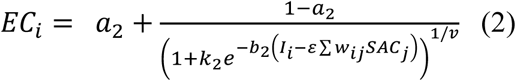

where *a*_*1*_=-0.1, *a*_*2*_=-0.05 were the activity levels’ lower bounds, *b*_*1*_= 70, *b*_*2*_*=*10, *v = 2*.*5* controlled the steepness of the sigmoids, 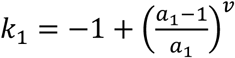 and 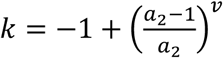 set the neurons’ activity to zero when the net input was zero, ε controlled the strength of interglomerular inhibition, and *w*_*ij*_ was the net connection strength from glomerulus *i* to glomerulus *j* as defined in the network construction above. The input *I*_*i*_ was the activity of the OSN connected to the *i*th glomerulus. The result was a system of 2*n* algebraic equations, where *n* was the number of glomeruli in the network, which we solved via MATLAB’s *fsolve* function.

### Model Inputs

To simulate biologically-constrained patterns of OSN input, we used published datasets consisting of matrices of OSN activity measured across many glomeruli in response to a large array of odorants (Ma et al., 2012; Burton et al., 2019). The first dataset, derived from Burton et al. (2019) (Fig. 3A), contained responses of OSN terminals in 105 glomeruli to 71 odorants (eliminating 8 odorants that did not evoke a response in any OSNs). The second dataset, derived from Ma et al. (2012), contained the responses in 94 glomeruli to 59 odorants at three concentrations (several odorant/concentration pairs were initially removed because they did not evoke a response in any OSNs, Fig. 6A). For both datasets, all OSN responses were normalized by the largest value in the dataset so that all values were bounded between 0 and 1.

### Measures of EC Activity

We calculated the lifetime sparseness of ECs as a measure of the selectivity of responses of individual ECs (Fig. 4B, 6D). We used the formula from Davison and Katz (2007):

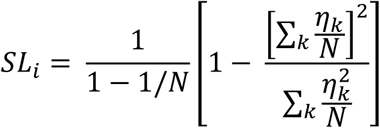

where *N* was the number of odorants presented to the network and *η*_*k*_ was the response of the EC of the *i*th glomerulus to odorant *k*. If an EC responded to only one odorant, the lifetime sparseness was maximal at 1 and if it responded to all odorants equally, the lifetime sparseness was 0. Similarly, we studied the sparseness of the patterns of activity evoked by odorants by calculating the number of ECs that were excited (activity above 0.045) or suppressed (activity below −0.07) by each odorant.

To examine the relationship between glomerular excitation and the prevalence of suppression, for each odorant we calculated the fraction of non-excited ECs that were suppressed (the number of ECs whose activity went below −0.07 divided by the number whose activity was below 0.045) as a function of the total excitation elicited by the odorant (the summed activity of ECs whose activity exceeds 0.045, Fig. 4D, 6F, 8F). We fitted the responses of the selective and nonselective networks with the rational function

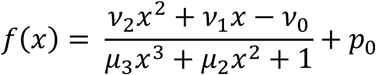

using MATLAB’s *fminsearch* function and the cost function

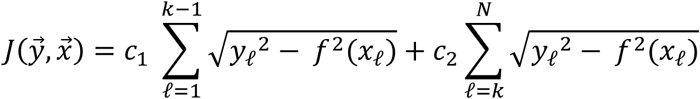

where *x*_ℓ_ was the total excitation of odorant ℓ and *y*_ℓ_ was the fraction of non-excited glomeruli that were suppressed by odorant ℓ, *k* was chosen such that *x*_*k*_ occurred near where the data saturates, *N* was the total number of odorants, and *c*_1_ > *c*_2_. Similarly, responses of the global networks that exceeded a threshold were fit using *fminsearch* and a quadratic polynomial

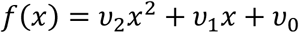

and the cost function

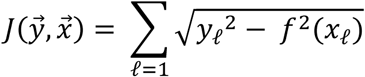

where *x*_ℓ_ and *y*_ℓ_ are the same as above. Below the threshold, the responses were fitted with a straight line.

We computed the Pearson correlation coefficient between odorant representation pairs using only OSNs/ECs that were responsive to at least one of the odorants. The decorrelation between the OSN and EC representations of odorant pairs was quantified using:

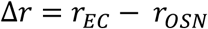

where *r*_*EC*_ and *r*_*OSN*_ were the Pearson correlation coefficients of the EC and OSN representations of the same odorant pair (Fig. 5, 8G).

### Interpolation Across Concentrations

To facilitate visualization of network effects on concentration-response functions (Figs. 7A, B), we expanded the OSN responses of the Ma et al. (2012) dataset by linearly interpolating OSN responses to concentrations between the three experimentally measured concentrations and linearly extrapolating responses to concentrations lower than those measured experimentally. For each odorant, we interpolated the response of every OSN using 99 artificial concentrations equally spaced between the lowest and middle concentrations and 99 artificial concentrations equally spaced between the middle and highest concentrations. If an OSN was not active at a measured concentration, an artificial concentration in an interval around the measured concentration was selected for it to become activated such that OSNs that were the most active at the next measured concentration were activated at the lowest concentration. To explore how odorants were represented at the lowest concentrations detectable by the olfactory system, OSNs responses that were nonzero at the lowest measured concentration were linearly extrapolated 100 steps (of the same size as those used for the interpolation above) to lower concentrations.

### Discriminability

We calculated the cosine distance between pairs of OSN and EC odorant representations using the mformula

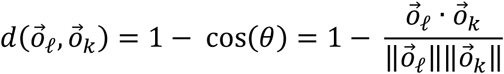

where 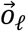 and 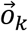 were column vectors with the activity levels of the neurons (either OSNs or ECs) representing odorant ℓ and *k* respectively and 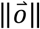 was the magnitude of the vector (Fig. 7C,E,F,G,H).

We modelled odorant representations of randomly distributed active neurons (either OSNs or ECs) using vectors 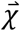 and 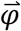 of length *d* with *p* and *q* active cells, represented by 1’s, respectively (Fig. 7E,F,G,H). We calculated that the expected cosine distance between 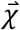 and 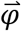 was:

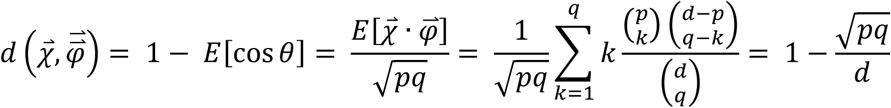

The result is a surface over a coordinate plane in which the height of the plane over the location (*p,q*) was the expected cosine distance between odorant representations with *p* and *q* cells active respectively. We compared this theoretical cosine distance to the cosine distances between odorant representations of our model inputs and outputs by plotting their cosine distances as a function of the number of active cells, averaging the cosine distance if multiple odorant pairs had the same number of active cells.

### Input-Tuned Networks

To examine the impact of input-tuned interglomerular networks (e.g., Fig. 8), we modified the network model described above such that the target sets of each glomerulus were chosen such that glomeruli inhibited those who had similar OSN input profiles. To do this, we created sets of artificial OSN inputs which were divided into four groups that preferentially activated different subsets of OSNs (Fig. 8A). The likelihood that each of the groups activated specific glomeruli was determined by four normal distributions (σ=8.5, truncated at 2σ) whose means were equally spaced across the arbitrarily indexed glomeruli. For each of the four distributions in an input set, we generated 71 OSN inputs with similar activity levels to 71 odorants from Burton et al. (2019) and with correlations spanning the full range of −1 to 1. We determined the target set of each glomerulus first by calculating the cosine distance between all pairs of OSN response profiles using the formula:

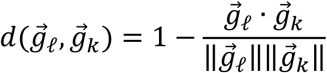

where 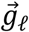 and 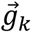 were column vectors of OSN ℓ and *k*’s response to all odorants. The target set of each glomerulus was then selected to be the 20 glomeruli whose response profiles had the smallest cosine distance. Once each glomerulus’ target was nonrandomly chosen, SAC connections were assigned as described above (Fig. 8B).

## Results

### Model Interglomerular Network Construction

We sought to determine whether broadly-distributed inhibitory connections, such as those mediated by SACs in the mouse OB (Aungst et al., 2003; Kiyokage et al., 2010; Banerjee et al., 2015), can mediate heterogeneous patterns of inhibition across a network with odorant-evoked sensory inputs to OB glomeruli. We also sought to explore how the structure of inhibitory connections between glomeruli impacts the input-output transformation of the OB network. We generated a reduced model of glomerular OB circuitry consisting of an array of approximately 100 glomeruli, each receiving OSN input and having a single excitatory output, EC, as well as a population of inhibitory interneurons representing SACs that mediate inhibitory connections to other glomeruli in the network (Fig. 1A). OSN inputs mediated feedforward excitation to the EC and SAC, and the EC in turn excited SAC associated with its glomerulus (thus, SACs received both mono- and disynaptic excitation); SACs mediated lateral inhibition through inhibitory connections onto ECs of other glomeruli (Fig. 1B). EC excitation (or inhibition) was used as a measure of glomerular output. Thus, in our simplified model, ECs encapsulate both mitral/tufted (M/T) cells, which are the principal OB output neurons, as well as external tufted cells, which mediate feedforward excitation to M/T cells and SACs (De Saint Jan et al., 2009; Gire et al., 2012; Banerjee et al., 2015; Liu et al., 2016). We chose this simplified structure, rather than attempting to recapitulate the entire glomerular circuit, because external tufted cells are thought to play a major role in driving M/T cell excitation (Hayar et al., 2004; De Saint Jan et al., 2009; Liu et al., 2013), and by merging the synaptic transfer functions of this feedforward excitatory pathway we significantly reduced the number of parameters and equations in the model. We made this transfer function highly nonlinear (Fig. 1C) to account for the very nonlinear responses of ET cells, which respond with spike burst to levels of sensory input above a certain threshold (Hayar et al., 2004; Gire and Schoppa, 2009). Although not explicitly included, the feedforward excitation was factored into the nonlinearities in the input-output functions of the EC and SAC (Fig. 1C). The result was a minimal network that was computationally efficient and tunable, but with its key features (i.e., combinatorial patterns of sensory input to glomeruli and SAC interglomerular innervation) constrained by experimental data. This minimal network allowed us to focus specifically on response properties that were determined by the structure of interglomerular inhibition, decoupled from the detailed cellular properties of individual neuron types and from intraglomerular or feedback inhibition mediated by other OB circuit elements.

We used several key findings from Kiyokage et al. (2010) to guide construction of the interglomerular network model. First, the SAC population included two morphologically distinct subtypes, termed oligoglomular and polyglomerular by Kiyokage et al. (2010), with differential abundance within the population. The “oligoglomerular” SAC subpopulation targeted relatively fewer glomeruli than the “polyglomerular” subpopulation but made up about 80% of the total population (See Methods for details). Second, the percentage of the total connection weight made by an SAC (and thus the inferred strength of inhibitory connection) was nonuniform across the glomeruli innervated by a particular SAC, but instead decayed roughly exponentially. Formally, the interglomerular connections of each SAC were wired to *h*_*o*_ or *h*_*p*_ glomeruli (representing oligoglomerular and polyglomerular SACs, respectively), chosen randomly and without repetition, each with a weight drawn from an exponential distribution modeling the percentage of SACs’ processes in each glomerulus in Kiyokage et al. (2010). This distribution is summarized in Figure 1D, where the weights of outgoing connections of a single representative SAC are shown in descending order. Finally, each glomerulus was associated with 40 SACs, based on previously-published estimates (Parrish-Aungst et al., 2007). As a result, the population of all SACs associated with a glomerulus - which we subsequently refer to as a SAC-glomerulus module - innervated a “target set” of many or relatively few other glomeruli, depending on the degree to which the different ‘sister’ SACs of a glomerulus innervated the same or different glomeruli.

We used the degree of selectivity of this glomerular targeting, defined by the “target set” of *m* glomeruli innervated by one SAC-glomerular module, as the primary independent parameter for exploring how interglomerular network structure impacts olfactory processing. All connections sent by a single SAC-glomerulus module can only connect to nodes in the target set *m*. The total strength of connection from one module to another is then the sum of the weights of all the SAC connections originating from that glomerulus. Varying the parameter *m*, the size of the target set of glomeruli, produced “selective” networks with fewer, stronger connections when *m* was low, or “nonselective” networks with more and weaker connections when *m* was high.

The connectivity matrix of a representative network of each type is shown in Figure 1E. Note that the color scale bar is the same for both figures, and the difference in color represents the overall difference in connection strengths.We also considered “globally” connected networks as explored in earlier theoretical work (Cleland and Sethupathy, 2006; Polese et al., 2014), in which each SAC-glomerulus modulewas connected to all other glomeruli with equal strength. Tthe same average strength of connections was set to be the same as in the selective and nonselective networks (see Methods for details). Thus, in the subsequent figures and text we refer to “selective” (*m*=20), “nonselective” (*m* equal to 94 or 105, the total number of glomeruli in the network) or “global” networks.

To model neural activity in the cells we used a firing-rate-type model in which two families of algebraic equations were associated with ECs and SACs (Fig. 1C) that modeled their deviation from spontaneous firing rates. Note that because the plots represent response relative to spontaneous firing rate the negative part of the input-output function corresponds to the suppression of spontaneous action potential firing. The shallower slope of the SAC response function, relative to that of the EC, was chosen based on recent reports of a log-linear relationship between odorant concentration and SAC activation *in vivo*, as measured by calcium indicator fluorescence changes (Banerjee et al., 2015).

### SAC networks can support heterogeneous interglomerular inhibition

We first examined how the strength of interglomerular connections formed by a single SAC-glomerulus module is affected by the selectivity of interglomerular inhibition (represented by the size of the target set, *m*) by examining only the connectivity matrices of the interglomerular networks. We measured the total strength of each module’s outgoing connection to another glomerulus by summing the weight of all connections from the source glomerulus to the target glomerulus. Figure 2A shows the strength of all outgoing connections of a representative SAC-glomerulus module to the 93 other glomeruli in an example network with *m*=20 (selective, top), *m* = 93 (nonselective, middle) and global (bottom) networks. When *m*=20, the glomerulus did not connect to the vast majority of the other glomeruli in the network, although the few connections it did form were very strong. As *m* increases, the connection strength became much more uniform. When connectivity was global, all the glomeruli received connections and the strengths of these connections was homogeneous and generally weaker than the *m* = 93 case. These observations from a single representative glomerulus are generalized in Figure 2B, which shows the distribution of outgoing connectivity strength from 100 simulations of networks with selective (*m* = 20, top), nonselective (*m* = 93, middle), or global (bottom) connectivity. When *m* = 20, the distribution was very wide with large zero strength probability. As *m* increased the distribution shifted to the left and became much narrower with a smaller zero strength probability. Overall, when inhibition was selective, interglomerular connections were heterogeneous with the few connections that are formed having a high average strength as well as highly variable strength. Conversely, as *m* increased (i.e. selectivity decreased) interglomerular connections became more homogeneous as well as weaker.

**Figure 2.**
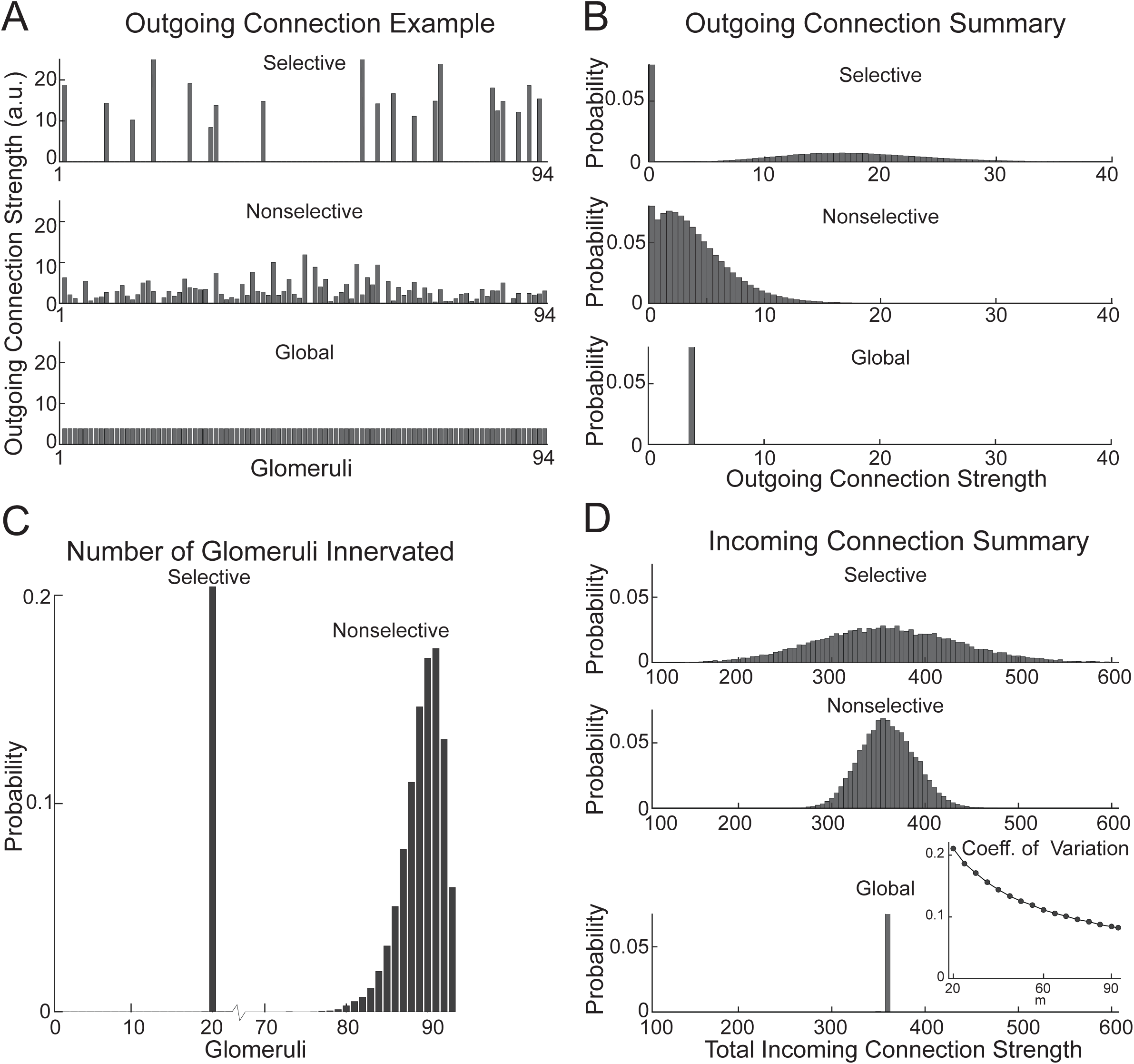
SAC Mediated Interglomerular Connectivity is Heterogeneous. **A**. Examples of the strength of outgoing connections formed by a single glomerulus in selective, nonselective, and globally connected networks. **B**. Distributions of the strengths of outgoing connections made by all glomeruli. **C**. The distributions of the number of glomeruli innervated by a single glomerulus in selective and nonselective networks. **D**. Distributions of the total incoming strength of connections received by glomeruli. Inset: The coefficient of variation of the total incoming strength as a function of selectivity (*m*).

We next examined the extent of connectivity as a function of selectivity of inhibition (Fig. 2C). When *m* was relatively small, each SAC-glomerulus module tended to connect to every glomerulus in their target set. As *m* increased, the fraction of the target set receiving connections decreased. Note that, although possible, it was very unlikely that a single glomerulus would connect to every other glomerulus in the network even when *m* = 93. We also examined the distribution of the strength of incoming connections that each glomerulus received across different values of *m* (Fig. 2D). The mean of the distribution was constant because the total number of SAC connections made was constant as *m* was varied, but the distributions narrowed as *m* increased. The coefficient of variation of the distributions of incoming connection strength decreased as *m* increased (Fig. 2E), indicating that the inhibition received by each glomerulus was much more variable in the sparse, *m* = 20, condition compared to the broad, *m* = 93, condition. These results demonstrate that, despite a high divergence of SACs from one glomerulus to others and despite the fact that multiple such SACs extend from each glomerulus, inhibitory networks mediated by SACs can, in theory, result in heterogeneous patterns of inhibition across the glomerular array. The degree of heterogeneity depends on the selectivity of this interglomerular connectivity. Our results also confirm that changing *m* widens the distribution of interglomerular inhibition without affecting the average inhibition between glomeruli, allowing us to compare the effect of the breadth of SAC connectivity in addition to the effect of the strength of inhibition on the patterns of EC suppression and excitation elicited by OSN inputs.

### Effect of SAC network structure on glomerular outputs

We tested the effect of selectivity in SAC connectivity on the transformations of OSN inputs performed by this simplified network. OSN inputs were taken from previously-published data consisting of optical signals representing activity of OSN populations converging onto distinct glomeruli, in response to a large panel of odorants (Ma et al., 2012; Burton et al., 2019). The initial dataset included 71 odorants across 9 chemical classes, tested at a single concentration and imaged across 105 glomeruli in a single field of view (Burton et al., 2019). Response patterns were converted into magnitudes of input to each glomerulus in the model array, normalized to the strongest observed OSN response (Fig. 3A). Different odorants evoked different patterns of OSN input across the glomerular array and covered a range of sparseness values. Across all input patterns, there was an approximately linear relationship between OSN input strength and total amount of SAC activation (Fig. 3F), consistent with previously-reported experimental results (Banerjee et al., 2015).

**Figure 3.**
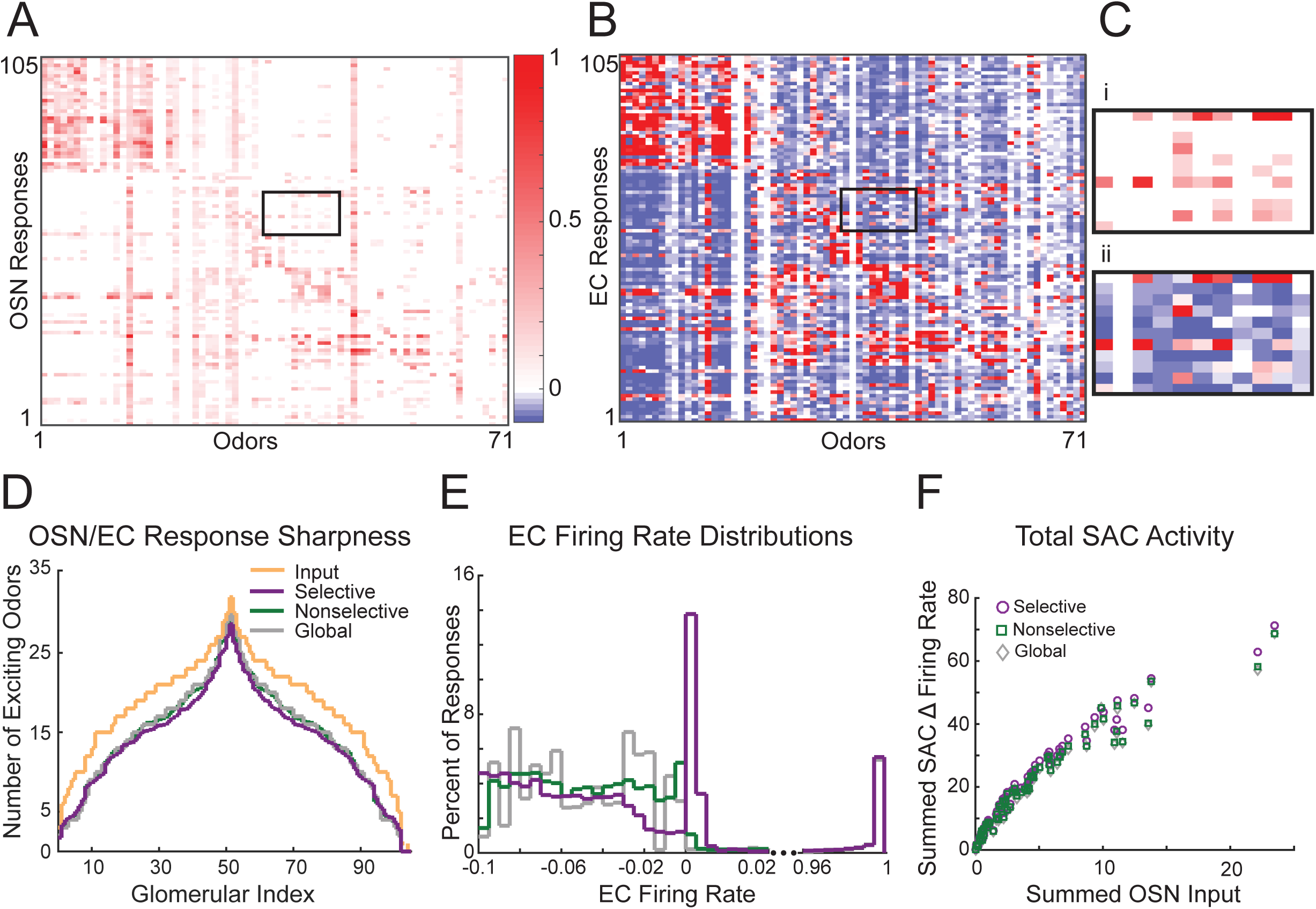
SACs Mediate Sparse, Heterogeneous Inhibition. **A**. Experimentally measured OSN responses to the 105 odors from Burton et al. (2019). **B**. EC responses of a selective network to the odors in A. **C**. A subsample of the OSN responses (i) and corresponding EC responses (ii) in A and B. The locations of the subsample are indicated by black boxes in A and B. **D**. The number of exciting odors per OSN glomerulus for selective, nonselective and global networks. **E**. Distribution of EC firing rate responses to the 105 odors from Burton et al. 2019, for each network class. Only responses near 0 (suppressive and week excitatory) and near 1 (strong excitatory) are shown. **F**. The summed changed in firing rate of SACs (averaged across networks) as a function of summed OSN input for each odorant.

Next, the system of algebraic equations representing the glomerular network was solved (see Methods), yielding an array of output values for the EC from each glomerulus. The EC response to an odorant was bounded between −0.1 and 1 and was classified as “excited” if it exceeded an upper threshold of 0.045, “suppressed” if it was below a lower threshold of −0.07, or “neutral” between these two values. We assessed the impact of network connectivity on glomerular outputs by varying selectivity values (*m*) and mean inhibitory strength (*ε*). For each parameter setting, we measured the distribution of EC activation by generating 50 networks and calculating their responses to all odorants (an example network response to a collection of input odorants is shown in Fig. 3B).

The primary transformation mediated by the SAC network was, as expected, the introduction of suppressive responses that were not present at the input stage. In addition to suppressing EC responses below baseline, the inhibitory SAC network also transformed input representations by eliminating weakly excitatory responses, resulting in a sparsening of the glomerular response map (light red indices in Fig. 3Ci compared to Fig. 3Cii). All networks, regardless of selectivity or inhibition strength, sharpened glomerular tuning, with EC outputs from individual glomeruli activated by fewer odorants than their corresponding OSN inputs (Fig. 3D). All networks also produced a variety of suppressive responses in EC (Fig. 3E).

We separately examined the impact of selectivity and inhibition strength on network output, focusing first on how these parameters impacted the relative proportions of excited, suppressed and neutral glomeruli. Both excitation and suppression are fundamental measures of neuronal networks known to affect their function. As expected, these proportions were strongly impacted by the strength of SAC inhibition, with network output shifting from being dominated by excitatory outputs to being dominated by suppressed outputs as inhibition strength increased (Fig. 4A). Numbers of excited and suppressed glomeruli were roughly equal at ε values of 0.001 (Fig. 4Aii). Notably, inhibitory network selectivity (*m*) had little impact on this balance at a given ε value, at least when averaged across all odorant input patterns (Fig. 4A).

**Figure 4.**
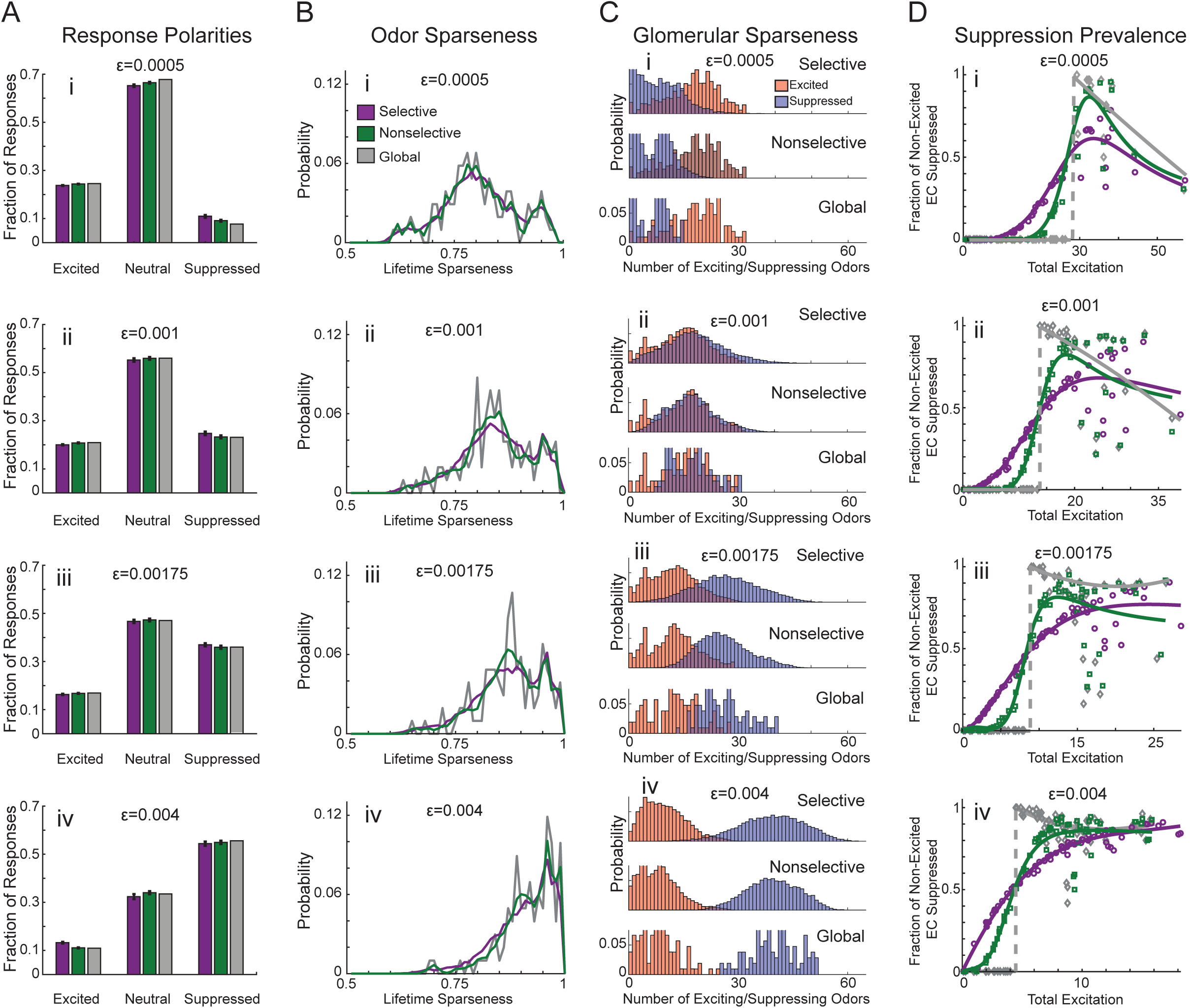
Excitation-Suppression Relationships Distinguish Global Inhibitory Networks from Selective and Nonselective Networks. **A**. The fraction of EC responses classified as excited (exceeding 0.045), suppressed (going below −0.07), or neutral (between −0.07 and 0.045). Selective networks are purple, nonselective are green, and global are gray. Rows (i-iv) differ in inhibition strength (ε) as indicated. **B**. Distributions of lifetime sparseness of EC responses. **C**. Distributions of the number of odors eliciting excitation and suppression in ECs. **D**. The probability of suppression occurring in ECs that were not excited, plotted as a function of summed excited EC activity. Each point is the mean of a single odor’s responses across networks. Selective and nonselective network fits are done with a rational polynomial (see Methods for details); the global network fit is done using a straight line followed by a quadratic polynomial.

To characterize the impact of network structure on individual odorant representations, we computed the lifetime sparseness of the network responses to all odorants (see Methods) (Fig. 4B). Similarly, we characterized the network impact on the representation of individual odorants by computing the number of excitatory and suppressive responses elicited by each odorant (Fig. 4C). Increasing inhibition strength (ε) led to an overall increase in sparseness of odorant responses (Fig. 4B), and an increase in the number of odorants eliciting suppressive responses in a glomerulus (Fig. 4C). Intermediate values of inhibition strength led to roughly similar distributions for the number of exciting or suppressing odorants per glomerulus (Fig. 4C). Again, the selectivity of the SAC network had little impact on the distributions of the number of exciting or suppressing odorants and lifetime sparseness. These results suggest that the selectivity of the SAC-mediated interglomerular network does not play a major role in sharpening odorant representations with respect to response spectra of individual glomeruli or sparseness of glomerular representations.

Finally, we examined the relationship between the prevalence of suppressed and excited glomeruli within a single odorant response. With lateral inhibitory networks, suppression of non-excited glomeruli should emerge and scale as the total amount of excitation to the network increases; this relationship is thought to mediate an intensity-dependent gain control of glomerular output (Cleland and Sethupathy, 2006; Banerjee et al., 2015). Here, we found that this function depended heavily on the mean strength of inhibition in the network. At the weakest inhibition strength (ε=0.0005), the prevalence of suppressive responses did not scale monotonically with total excitation for any network structure (Fig. 4Di). Instead, increases in total excitation beyond a certain value led to a reduction in the fraction of suppressed glomeruli, as excitation outweighed interglomerular inhibition (note the low values of all fitting curves in Fig. 4Di when excitation is high). Doubling the mean inhibition strength between glomeruli (ε=0.001) reduced this effect, with suppressive responses persisting at the highest levels of excitation (Fig. 4Dii), and further increases in inhibition strength (ε=0.00175 and ε=0.004, Fig. 4Diii and 4Div, respectively) led to excitation-suppression relationships that were closer to monotonic as net excitation levels increased across the glomerular array.

Notably, however, the structure of the interglomerular SAC network had a substantial impact on the relationship between excitation and suppression. In particular, at a given inhibition strength, less selective networks led to increasingly nonlinear relationships between total excitation and prevalence of suppression. For example, at the two highest strengths of inhibition, selective SAC networks resulted in a gradual, near-linear increase in the prevalence of suppressed glomeruli as total excitation increased, saturating at the highest levels of excitation (purple plot, Fig. 4Diii and 4Div), while nonselective networks resulted in a steeper relationship better fit by a sigmoid (green plot, Fig. 4Diii and 4Div). Finally, networks with global connectivity yielded a discontinuous relationship between total excitation and suppression at all inhibition strengths, in which no glomeruli are suppressed when total excitation is low, but when excitation exceeds a certain threshold, nearly all non-excited glomeruli become suppressed (gray plots, Fig. 4D). This discontinuity resulted from the uniform strength of inhibition between all glomeruli: when total excitation is low, all glomeruli are inhibited uniformly and weakly such that even glomeruli receiving no OSN input are not sufficiently inhibited to be considered suppressed; however, as total excitation increases, the strength of global inhibition becomes sufficiently strong to suppress most or all non-excited glomeruli. (This relationship persists even if 10% variability is added to the strength of the global connections to break the symmetry, and it also persists if the input-output curve of the EC cells shown in Fig. 1C is made less steep, similar to the steepness of the SAC curve of Fig. 1C). Overall, these results suggest that SAC networks with selective, rather than global, inhibition between glomeruli, allow for a more linear relationship between total input strength and prevalence of suppressive odorant responses, and are better able to match recent experimental data (Banerjee et al., 2015; Economo et al., 2016).

### Network Function

Next, we explored how SAC network structure alters the representation of odorant identity by the glomerular array. Previous experimental work has implicated the SAC network in decorrelating odorant representations by M/T cells (Banerjee et al., 2015), such that the correlation coefficient between pairs of odorant representations is lower at the level of glomerular output than at the level of OSNs (see Methods). To evaluate the impact of network structure and strength on input-output transformations with our model, we measured the correlations of EC odorant responses to odorant pairs resulting from selective, nonselective, and global networks.

To illustrate, the impact of the SAC network on the correlation between two representative pairs of odorants is shown in Figure 5A. The correlation coefficient for the OSN input patterns for each pair is shown by the dashed line (*r* =0.49 and 0.83 for the upper and lower pair, respectively). Each plot shows the distribution of correlation coefficients for the EC output patterns across 100 independent realizations of either selective (*m*=20, purple) or nonselective (*m*=93, green) networks. All iterations of both selective and nonselective networks led to lower correlation coefficients, indicating a decorrelation of the glomerular odorant representations for each odorant pair, consistent with previous models of OB network function (Cleland and Sethupathy, 2006; Wiechert et al., 2010; Banerjee et al., 2015). In addition, the selective networks yielded a wider range of correlation coefficients across the 100 network iterations in both cases (i.e., the purple distribution is wider than the green, Fig. 5A).

**Figure 5.**
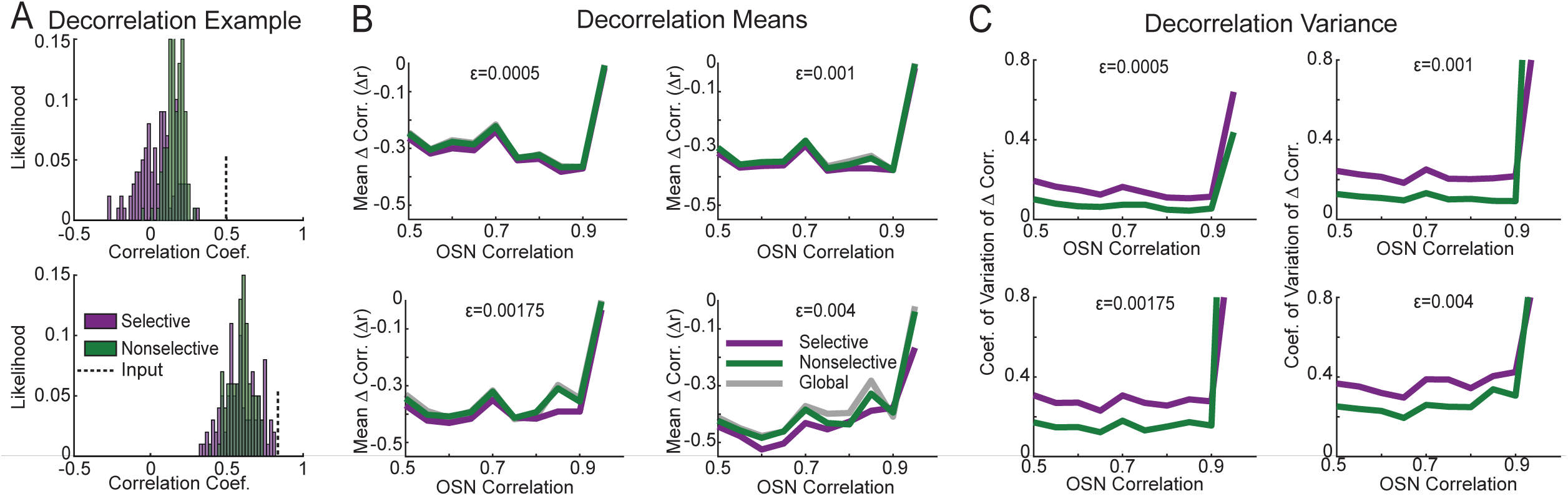
Lateral Inhibition Decorrelates Odor Representations. **A**. Examples of the distribution of correlation coefficients responses to two odor pairs over 100 realizations of selective and nonselective networks. Dashed lines indicate the correlation coefficient of the OSN odor representations. **B**. Mean decorrelation over 100 network realizations of odor pairs (*r*_*EC*_ − *r*_*OSN*_)plotted as a function of their OSN correlation. **C**. Coefficient of variation of odor pairs’ decorrelation across 100 selective and nonselective network realizations, plotted as a function of their OSN correlations. Only a single global network was created for each ε so no coefficient of variation could be calculated.

To expand this analysis to all possible odorant pairs, we computed the input (OSN) correlation coefficient for each pair, and the mean correlation coefficient of the EC responses to the same pair across the 100 randomly generated network iterations. We then calculated the amount of decorrelation by the SAC network (i.e., the reduction in correlation coefficient from OSN input to EC output) as a function of the OSN input correlation, for all odorant pairs with input correlation above 0.5 (Fig. 5B). All network structures decorrelated inputs to a similar degree (Fig. 5B), with the exception of the strongest level of inhibition (ε=0.004), where selective networks generated modestly greater decorrelation of OSN input patterns than nonselective or globally-connected networks. In addition, all networks decorrelated highly correlated odorant pairs (*r*_*OSN*_>0.95) significantly less than moderately correlated pairs (*r*_*OSN*_<0.95). Decorrelation in moderately correlated odorant pairs results from ECs in a particular glomerulus receiving enough net excitatory input to become excited by one odorant of a pair but not the other. In contrast, odorant pairs with highly correlated OSN representations evoked activity that was so similar that the ECs of few or no glomeruli were excited by only of the odorants. This resulted in highly correlated activity and low decorrelation (Fig. 5B).

Importantly, the plots in Figure 5B show the mean degree of decorrelation across all 100 instances of each network structure. However, as seen in Figure 5A, this value could vary considerably for different instances of a network. Variability arose due to the heterogeneous connectivity between different glomeruli, and the fact that each odorant activated a different combination of glomeruli. Thus, the degree of decorrelation varied with the odorant pair and, for a given odorant pair, with the particular glomerular connectivity at each network instance. Figure 5C shows this variation, plotted as coefficient of variation of decorrelation across the 100 network instances (global networks did not have a coefficient of variation because only a single network was created). Notably, the amount of this variation in decorrelation was greater for selective networks than for nonselective networks (Fig. 5C). In addition, this variability increased with the mean strength of inhibition (ε) in the network, particularly for selective networks and particularly for odorant pairs with a high OSN input correlation. These results indicate that while all connectivity schemes decorrelate odorants similarly on average, particular instances of selective networks have the capacity to decorrelate odorant representations significantly more than unselective or globally-connected SAC networks.

### Odorant Concentration and Discriminability

Earlier computational and experimental studies have suggested that interglomerular inhibition mediated by the SAC network contributes to concentration-invariant odorant representations by implementing an intensity-dependent gain control across all glomerular inputs (Cleland and Sethupathy, 2006; Banerjee et al., 2015). These predictions arise from assumptions of global (or near-global) SAC connectivity. Thus, we next sought to examine the effect of SAC network structure on input-output transformations as a function of odorant concentration. To test this using OSN input patterns constrained by experimental data, we used a different previously-published dataset that included OSN inputs to 94 glomeruli imaged in response to three concentrations of the same odorants (Ma et al., 2012) (Fig. 6A).

**Figure 6.**
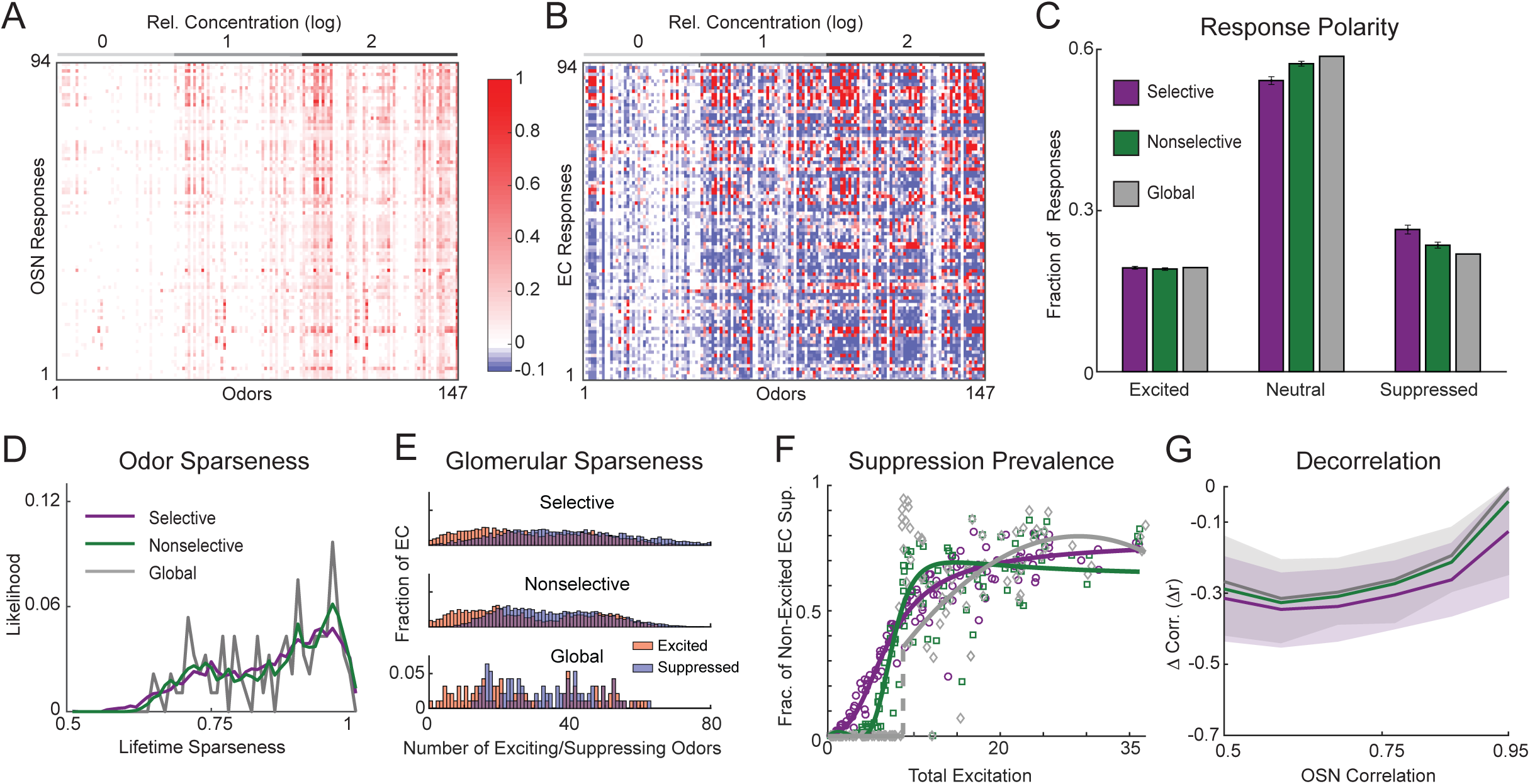
Networks Reproduce Key Responses and Computations When Driven by Alternative, Experimentally Derived OSN Inputs. **A**. Experimentally measured responses to odors from Ma et al. (2012). Most odors are repeated at 3 concentrations. Concentration is indicated above the panel. **B**. EC responses of a selective network (ε=0.00175) to the odors in A. **C**. The fraction of EC responses that are classified as excited (exceeding 0.045), suppressed (going below −0.07), or neutral (between −0.07 and 0.045) across networks of each type with ε=0.00175. **D**. Distributions of the lifetime sparseness of ECs. **E**. Distributions of the number of odors eliciting excitation and suppression in ECs. **F**. The probability of suppression occurring in ECs that were not excited, plotted as a function of summed excited EC activity. Each point is the mean of a single odor’s response across networks. Selective and nonselective fits are done with a rational polynomial and the global fit is straight line followed by a quadratic polynomial. **G**. Median decorrelation of odor pairs from their OSN to EC representations (*r*_*EC*_ − *r*_*OSN*_, solid lines) plotted as a function of OSN correlations. The first and third quartiles of the distributions for the global and selective networks form the shaded areas.

First, we confirmed that our SAC network model behaved similarly with this input dataset as with our initial dataset. As in the initial characterization, we measured the distribution of EC activations in response to each odorant response in the Ma et al. (2012) dataset across all three connectivity schemes (see Methods). An example EC response matrix of a selective network with intermediate connection strength (ε=0.00175) is shown in Figure 6B. We found similar effects of SAC network inhibition strength and network structure with this new input dataset. For example, the strength of inhibition in the network had similar effects on the lifetime sparseness and relative prevalence of excited and suppressed glomeruli, with a similar balance at intermediate inhibition strengths (ε=0.00175 or 0.001), and with network structure having little impact on these proportions (Fig. 6C,D,E). As with the Burton et al. (2019) dataset, network structure did impact the relationship between total excitation and fraction of suppressed glomeruli, with the most linear relationship occurring with selective networks and a nonlinear relationship occurring with the globally-connected network (Fig. 6F, compare with Fig. 4E). Finally, all SAC network structures decorrelated OSN input representations to a similar degree (Fig. 6G). These results suggest that the effects of the SAC network model on input-output transformations across the glomerular array are generalizable across different experimental datasets.

Next, we investigated intensity-dependent transformations by the network by comparing concentration-response functions for inputs and model EC outputs from individual glomeruli as well as across the population. To facilitate the comparison of OSN and EC concentration-response functions, we generated simulated OSN responses at concentrations not included in the original dataset by interpolating responses for each OSN between the three experimentally-tested concentrations and, in some cases, extrapolating to fill in concentrations lower than those measured experimentally (see Methods). Figure 7A (left panel) shows an example of OSN responses to a single odorant across concentrations. The remaining panels of Figure 7A show the corresponding EC activity evoked in networks with each connectivity scheme. The threshold concentration for the emergence of suppressive responses among ECs is highly variable in the selective and non-selective networks, while in the global network it is stereotypical, with all ECs showing suppression at the same concentration. This property underlies the discontinuity observed in the excitation-suppression relationship of global networks (e.g., Fig. 4D), and does not match experimental observations in which suppressive responses emerge at a range of concentrations (Banerjee et al., 2015; Economo et al., 2016). Variability in concentration-response functions across different network instances was largest among selective networks, because of their relatively greater variability in connectivity; this variability is highlighted in plots of the OSN inputs and EC outputs of three individual glomeruli across three realizations of each type of network (Fig. 7B).

**Figure 7.**
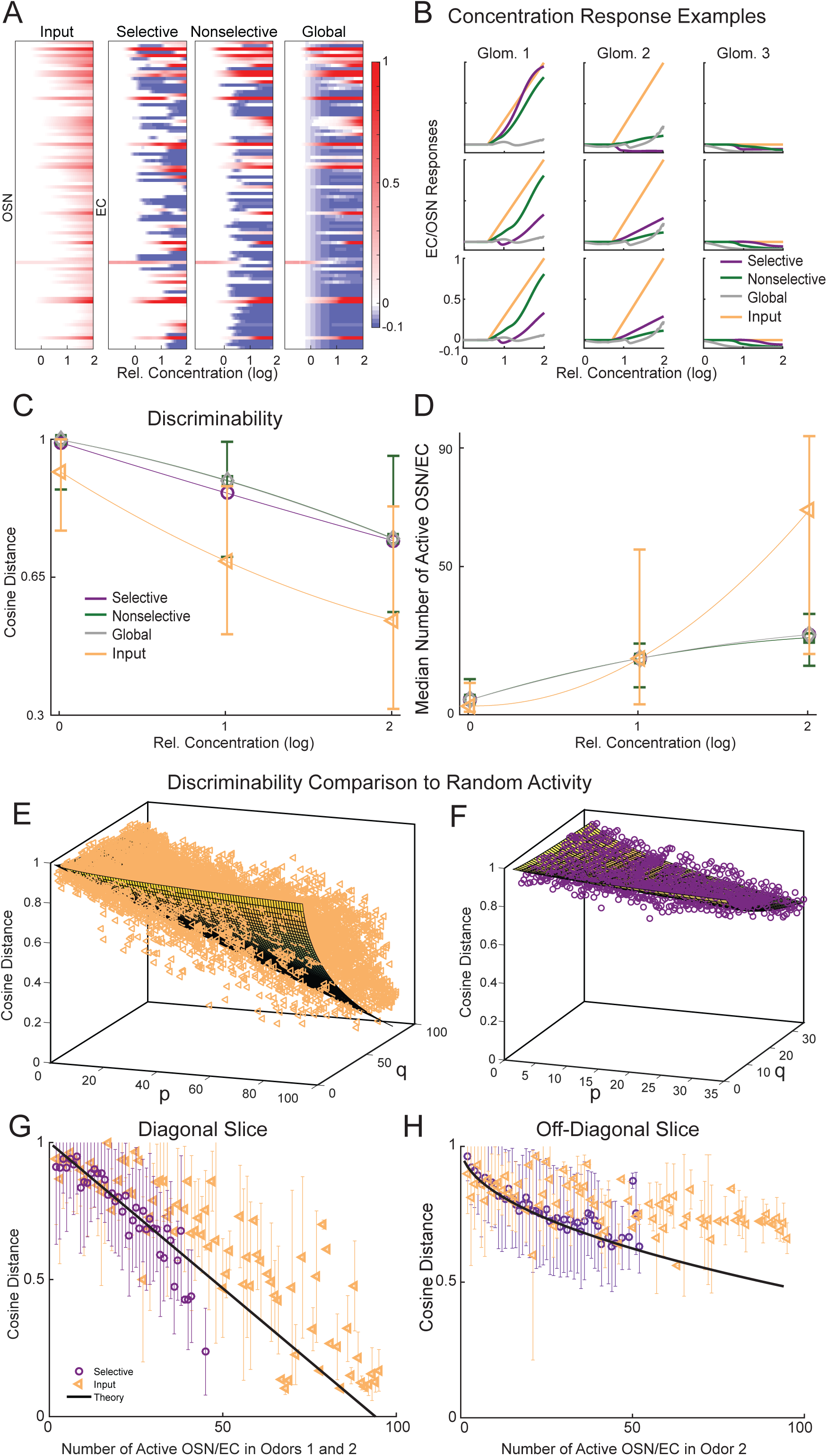
Lateral Inhibition Maintains Odor Discriminability Across Concentrations by Sparsening Representations. **A**. Left panel: OSN responses to a single odor over three orders of magnitude in concentration, generated from the Ma et al. dataset (see Methods). Right panels: EC firing rate responses (with ε=0.00175) to the odor in the left panel. **B**. Inputs and outputs of three glomeruli (columns i, ii, and iii) across three independent realizations of each network type (rows). **C**. Median cosine distances between odor representations both in the OSN and EC layers as a function of concentration. Error bars indicate quartiles of the OSN and nonselective networks’ distributions. Points are connected by quadratic fits for visualization. **D**. Median number of active OSNs and ECs as a function of concentration. Error bars indicate quartiles of the OSN and nonselective networks’ distributions. Points are connected by quadratic fits for visualization. **E**. Mean cosine distance between OSN representations of odors as a function of the number of active OSNs in the first (*p*) and second (*q*) odor respectively. Surface is a theoretical prediction of the mean cosine distance between odor representations with randomly activated cells (OSN or EC). **F**. Same as E with EC representations. **G**. A diagonal slice (*p*=*q* is the x axis) of the surface and data points in E and F. Error bars indicate the standard deviation of cosine distances across odorant representation pairs with the same number of active cells (some error bars are very small and hence difficult to distinguish from the markers). **H**. An off-diagonal slice (*p*=25, *q* is the x axis) of the surface and datapoints in E and F.

We then explored the effect of concentration on discriminability by computing the cosine distance between all pairs of OSN odorant representations at each of the three experimentally-measured concentrations, as well as the cosine distance between all EC odorant representation pairs, for each network connectivity type with ε=0.00175 (Fig. 7C, markers are medians, error bars are quartiles). As concentration increased, the cosine distance between OSN representation pairs decreased, indicating an increase in pattern similarity and a resulting decrease in discriminability. This effect was reduced among EC representations for all network types, indicating that interglomerular inhibition mitigated the loss of discriminability at higher concentrations that arises from increasing overlap in OSN input patterns.

It is important to note that the cosine distance is highest for pairs that are represented by non-overlapping sets of cells and that it is easier to maintain non-overlapping sets if fewer glomeruli are involved. Thus, the number of active glomeruli in each odorant representation may play an important role in discriminability. The number of active OSNs/ECs per odorant increased with concentration; the number of active OSN inputs grew rapidly (Fig. 7D), while the number of active EC outputs remained much lower due to interglomerular inhibition (yellow markers compared to purple/green/gray markers, Fig. 7D). The growth in the number of active glomeruli among both OSNs and ECs closely mirrored the drop in the cosine distance across concentration, suggesting that the number of active glomeruli might be the determining factor in odorant discriminability. Therefore, the ability of the OB network to maintain odorant discriminability across concentrations might be primarily determined simply by sparsening the response, as opposed to a transformation of response patterns that more efficiently encodes odorant identity by virtue of their particular correlation relationships.

To test this possibility, we analytically calculated the expected cosine distances of theoretical activity patterns in which the correlational structure in the patterns was eliminated and only response sparseness was represented (see below). We then compared this theoretical expected cosine distance - sparseness relationship to that taken from OSN or EC response patterns in the experimentally-derived datasets. If, indeed, solely the number of cells activated determines discriminability, the theoretical and experimentally-derived cosine distances should be similarly distributed. However, if changes to the correlational structure of EC responses plays a significant role in determining discriminability among output patterns, then the distribution of cosine distances between EC responses should deviate from the theoretical relationship.

To analytically compute the expected cosine distance between odorant representations with no correlation structure, we considered theoretical odorant representations whose number of active cells were within the same range as the experimental set. The identities of active cells were chosen randomly and independently, and their activity levels were set to 1. A formula describing the expected discriminability of odorant representation pairs (as measured by the cosine distance) was derived. For a pair of odorants that have *p* and *q* active cells, the expected cosine distance was

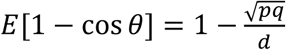

We generated a surface of this expected distance over the (*p,q*) plane (grid in Fig. 7E,F). If the cosine distances between the OSN or EC response patterns in the actual dataset were determined purely by the number (and not the identity or the relative firing rates) of the active glomeruli, then the model data should fall very close to the theoretical surface. We compared the theoretical surface of expected cosine distances of *p* and *q* randomly activated cells to the cosine distances of pairs of OSN or EC responses from a single selective network composed of *p* and *q* active cells (yellow triangles in Fig. 7E, purple circles in Fig. 7F, respectively). We also took two 2-dimensional vertical slices, with the line in these plots corresponding to a slice through the theoretical surface, and the symbols the same as those in the full surface plots, with error bars (st.dev.) added (Fig. 7G,H). The cosine distances of EC representations of odorants fit the theoretical expected value surface well, except for cases when both odorants have a high number of active glomeruli (Fig. 7F,G,H).

Interestingly, we found that the distances between OSN input representations did deviate from those predicted by the theoretical distanse-sparseness relationship, falling mostly above the theoretical surface/curves (Fig. 7E,G,H). In other words, the experimental OSN dataset had less overlap between odorant pairs than would be expected randomly based on the number of glomeruli activated, consistent with a nonrandom correlational structure in the odorant response specificities of OSN inputs to glomeruli. When comparing representations with the same number of active glomeruli, EC odorant-response pairs had smaller cosine distances and less discriminability relative to OSN pairs (yellow triangles are generally above the purple circles and black lines in Fig. 7G,H). However, across concentrations, EC representations were sufficiently sparse that they were much farther apart than their OSN counterparts (purple/green/gray lines are above the yellow line in Fig. 7C).

Overall, this analysis indicates that the increase in discriminability of EC odorant representations relative to OSN representations is driven primarily by reductions in the numbers of active glomeruli (i.e., sparsening) and not by changes to the correlational structure of their population activity patterns. It also raises the possibility that if inhibitory interglomerular connections were tuned to reflect the non-overlapping structure of OSN inputs, perhaps through preferential inhibition of glomeruli with similar input profiles, the SAC network could improve the discriminability of OB odorant representations through sparsening while preserving the non-overlapping structure already present in OSN representations. We explored this possibility in the following set of simulations.

### Input-Tuned Networks

Earlier studies have suggested that interglomerular inhibition may be nonrandom (Girardin et al., 2013; Economo et al., 2016; Mohamed et al., 2019) and further that interglomerular inhibition that is “input-tuned”, such that glomeruli with similar response profiles are more likely to connect, may enhance odorant discrimination (Linster et al., 2005; Girardin et al., 2013). Thus, we asked whether directing the selectivity of SAC connectivity based on input tuning would produce different results than random connectivity. To test this, we generated idealized datasets of OSN inputs in which glomerular inputs had well-defined relationships to one another in terms of response tuning and explored the impact of selective SAC networks with particular connectivities related to this input structure. Each of the 25 artificial input datasets consisted of four sets of artificial odorants that activated OSNs corresponding to one of four normal probability distributions (truncated at 2σ) whose means were evenly distributed across the population (Fig. 8A). Individual odorant responses were constructed by taking the 71 odorant responses from Burton et al., 2019 and randomly reassigning the index of nonzero responses using each of the four distributions (for a total of 284 odorant responses in each input dataset). We generated input-tuned selective networks by creating SAC target sets (*m* = 20) for each glomerulus comprised of those glomeruli that respond most similarly toeach other across the artificial odorant panels (right panel, Fig. 8B). We also generated random selective networks (with randomly selected target sets) as described earlier (left panel, Fig. 8B) as well as nonselective networks. We then compared the effects of selective, nonselective, global, and input-tuned networks at two values of ε to test for interactions between the mean and variance of inhibition between glomeruli. This strategy allowed us to evaluate input-output transformations across many instances of network connectivity and to compare differences in network function for input-tuned versus randomly-connected networks.

**Figure 8.**
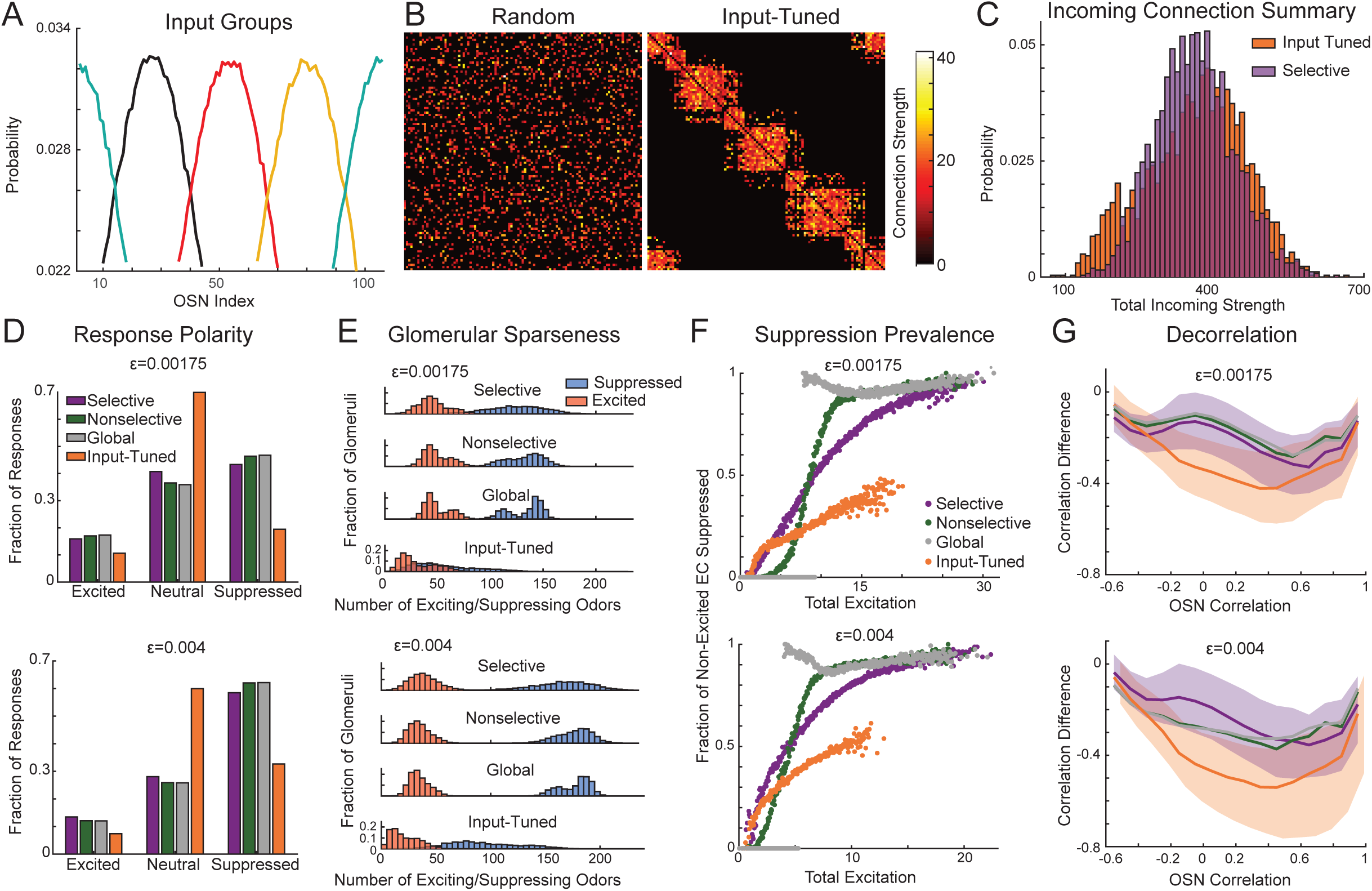
Input Tuned Networks Produce Improved Decorrelation. **A**. Distributions describing the probability that each of the 4 odor groups (represented by different colors) will activate each OSN. **B**. Adjacency matrices of random selective (left) and input-tuned (right) networks. **C**. Distributions of the total incoming strength of SAC connections for both random, selective (purple) and input-tuned (orange) networks. **D**.**-G**. Analysis of selective (purple), input-tuned (orange) and global (gray) network responses with ε=0.00175 (top panels) and ε=0.004 (bottom panels). **D**. The fraction of EC responses that are classified as excited, suppressed, or neutral. **E**. Distributions of the number of odors eliciting excitation and suppression in ECs. **F**. The probability of suppression occurring in ECs that were not excited, plotted a s a function of summed excited EC activity. For selective, nonselective, and input tuned networks, each point is the mean response of odors binned by total excitation. For global networks, only responses exceeding 0 suppressed glomeruli are binned in this way. **G**. Median decorrelation of odor pairs from their OSN to EC representations (r_EC_ − r_OSN_, solid lines) plotted as a function of OSN correlation. The first and third quartiles of the distributions for the functional and selective networks form the shaded areas.

These “input-tuned” networks transformed OSN input patterns in a manner distinct from that of selective but randomly-connected networks as well as the other connectivity schemes. First, the architecture of interglomerular connectivity differed from that of selective networks, with input-tuned networks exhibiting a broader distribution of the total strength of incoming SAC connections to a glomerulus (orange versus purple distributions, Fig. 8C). Second, input-tuned networks had four groups of glomeruli (corresponding to the four input distributions) with many reciprocal connections linked together by glomeruli that are in the intersection of two adjacent input distributions and consequently send and receive connections to and from glomeruli in adjacent groups. Thus, we found that input-tuned networks may differ from randomly connected networks both in straightforward topological measures like weighted in-degree distribution but also in more abstract topological measures like community structure, depending on the input statistics to which they are adapted.

Second, input-tuned SAC networks differed in their impact on OSN input patterns. Input-tuned networks resulted in sparser outputs than random networks, with the largest change being a decrease in the number of suppressive responses (Fig. 8D). Relatively more glomeruli responded with either excitation or suppression to a few odorants while not responding at all to the remaining odorants in the input-tuned networks, compared to random-selective and global networks (Fig. 8E). Thus, input-tuned network odorant representations were sparser and glomerular tuning sharper than in random networks. In addition, input-tuned networks maintained more balanced outputs and prevented the excessive suppression seen with greater mean inhibition in the selective and global networks when connection strength was moderately high (ε = 0.00175). The fraction of excitatory and suppressive responses elicited in the network when ε = 0.00175 was similar, particularly when compared to the responses in the random selective and global networks (Fig. 8D), and the distributions of the number of exciting and suppressing odorants per glomerulus were highly overlapping.

We next measured the relationship between total excitation and prevalence of suppression, as done with the responses to experimentally measured OSN input datasets in Figure 4E. Here, the random selective and nonselective networks and the globally-connected networks produced results similar to those seen with experimental inputs, with a linear-saturating increase in fraction of suppressed glomeruli with increasing total excitation in selective networks and a step-wise jump from no inhibition to near-total inhibition in globally-connected networks (Fig. 8F). Input-tuned networks produced a scaling relationship between excitation and suppression at either ε value tested, but with no saturation and a smaller amount of excitation overall. This further demonstrated that the structure of SAC networks substantially affects the relationship between excitation and inhibition.

Finally, we analyzed decorrelation of odorant representations by the three networks, calculated as in Figure 4. Here, we saw a substantial effect of network structure on decorrelation, with input-tuned networks decorrelating input patterns between median values of Δr_tuned_= −0.47 when r_OSN_=0 to its minimum of Δr_tuned_= −0.54 when r_OSN_=0.4, while the median decorrelation of random, selective networks varied between Δr_func_= −0.19 and − 0.33 over the same range (Fig. 8G, bottom panel, ε = 0.004). At the maximum amount of decorrelation (when the OSN correlation is between 0.2 and 0.6), input-tuned networks (with ε = 0.004) decorrelate odorant representations approximately twice as much as randomly and globally connected networks. This indicates that network connectivity determined by the statistics of the inputs, resulting in glomeruli preferentially inhibiting those that receive similar inputs, leads to an enhanced ability of the interglomerular network to separate overlapping odorant representations. Thus, adaptation of the interglomerular network to the statistics of stimuli presented to the network is important in producing balanced, decorrelated network responses.

## Discussion

We explored how the structure of a lateral inhibitory network affects computations in a simplified model of the mouse olfactory bulb. We constructed a mathematical model of SAC mediated lateral inhibition and glomerular responses to OSN input and considered four connectivity schemes: “selective”, where SACs in each glomerulus randomly connect among a random set of 20 target glomeruli; “nonselective”, where the set of possible target glomeruli is increased to include the entire network; “global”, where all SAC-glomerulus modules are connected to all others; and “input-tuned”, where SAC-glomerulus modules connect to glomeruli that receive similar OSN inputs. We found that selective and nonselective networks behave similarly with respect to many features of odorant representations. However, globally connected networks were distinct in that they produced no suppression when total excitation in the network was below a threshold and ubiquitous suppression of non-excited glomeruli when excitation exceeds a threshold, a result that is inconsistent with experimental data in which suppression scales with excitation (Economo et al., 2016). Finally, we found that organizing connectivity such that similarly tuned glomeruli are preferentially connected enhances decorrelation performed by the network.

Our model was constructed to be as simple as possible, taking into account only SAC anatomical data and including two functions found to be critical to OB activity: nonlinear feedforward excitation of output neurons and inhibitory interneurons and recurrent inhibition between glomeruli. By simplifying intraglomerular circuitry we were able to focus on the effects of interglomerular inhibition on responses to experimentally measured OSN input. The parameters of network structure were extrapolated from anatomical data from individual SACs (Kiyokage et al., 2010). We assumed that the synaptic efficacy between individual SACs and the glomeruli they innervate is governed by the percentage of their total processes innervating each glomerulus. In addition, we assumed that the strength of connection from glomerulus *i* to glomerulus *j* is the sum of all connections made by SAC in *i* innervating *j*. Thus, the percentage of the total SAC connection strength received by each glomerulus is similar to the percentage of total SAC processes entering each glomerulus observed in Kiyokage et al. (2010). Despite the detail in which SAC anatomy was modeled, our network framework is generic enough to incorporate additional interneuron anatomical data from the OB or to model another neural structure entirely.

We asked if interglomerular networks with data-driven connectivity properties and OSN input can produce heterogeneous and sparse inhibition, as has been observed in OB recordings in vivo (Soucy et al., 2009; Economo et al., 2016). We found that heterogeneity (as measured by the strength of connections) and sparseness (as measured by the number of glomeruli innervated) are intrinsic to SAC-mediated interglomerular networks, despite the broad interglomerular innervation of individual SACs. Although these features were enhanced in the selective networks with a broader distribution of connection strengths and many fewer outgoing connections, nonselective networks were also both heterogeneous and sparse compared to the globally connected networks. Notably, less than 10% of glomeruli in nonselective networks connect to all other glomeruli. Similarly, we find that the strength of interglomerular inhibition each glomerulus receives is heterogeneous. The distribution of incoming connection strength is most variable for selective networks but the coefficient of variation for nonselective networks is still approximately half that of the selective networks.

Although our network models have heterogeneous connections, it was not clear that they would produce selective suppression of glomerular output when driven by realistic OSN input patterns that activate multiple glomeruli. Examination of responses in selective networks showed that ECs receiving OSN input exhibited suppressed and neutral responses in addition to excited responses. Similarly, ECs that did not receive OSN input were suppressed to varying degrees or remained neutral. All networks produced a variety of suppressive responses. However, selective networks produced many more maximally suppressed responses (<-0.095) and, more importantly, more responses with little change in activity, than nonselective and global networks (∼0, Fig. 3E). Together with the connectivity calculations, this echoes previous experimental results in which suppression is combinatorial and glomerulus specific (Economo et al., 2016).

How do responses of selective, nonselective, and global networks differ? We found that many measures of activity, including the relative fraction of excited/neutral/suppressed responses, lifetime sparseness, and mean pattern decorrelation do not differ across connectivity schemes. However, global networks were distinguished from selective and nonselective networks by their relationships between the prevalence of excitation and inhibition. The fraction of suppressed ECs in selective and nonselective networks scaled with total excitation (for ε>0.001), matching experimental results (Economo et al., 2016). In contrast, global networks produced ubiquitous suppression when excitation exceeded a threshold and no suppression otherwise. These results indicate that, despite its use in numerous prior models of OB function (Cleland and Sethupathy, 2006; Polese et al., 2014; Banerjee et al., 2015) a globally connected network with uniform connection weights is unlikely to reflect the reality of interglomerular connectivity, at least in the mammalian OB.

Instead, we speculate that SACs constitute a network better described by our selective or nonselective models, although more investigation is necessary to distinguish between these two connectivity schemes. For example, both the total strength of incoming connection as well as the decorrelation of odorant representation pairs had similar mean values across randomly connected network structures. Experiments aimed at distinguishing between selective and nonselective connectivities could measure variability both in network structure and response to stimulation. Our result that selective and nonselective networks decorrelate odor representations equally well contrast with those from Wiechert et al. (2010) in which networks with sparse, strong connections decorrelated odorant pairs more effectively than those with many, weak connections. However, all of our network models were significantly less sparse than those used in Wiechert et al. (2010), which may account for the different results.

Similar to a previous study (Linster et al., 2005), we found that constructing interglomerular inhibition networks based on similarity of OSN input profiles improves contrast enhancement as measured by decorrelation between OSN and EC odorant representation pairs. Whether such input-tuned connectivity actually exists among glomeruli of the mouse OB remains unclear, although advances in our understanding of the structure of odorant coding space and its relationship to glomerular maps provide a platform for testing this hypothesis (Bozza et al., 2009; Chae et al., 2019). For example, inhibitory interglomerular connectivity could be domain specific, in which glomeruli receiving input from the same class (Bozza et al., 2009) preferentially inhibit each other and connections do not cross domain boundaries. Alternatively, inhibitory network structure may reflect odorant space with respect to behavioral meaning of the odorants. For example, recent work in the antennal lobe of the fly has found specific, non-random inhibitory connectivity between glomeruli involved in antagonistic behaviors (Mohamed et al. 2019).

Our results also support a role for recurrent inhibition and output normalization in maintaining dissimilar odor representations (for review, see Carandini and Heeger, 2012). Previous work demonstrated that purely feedforward inhibitory networks perform much worse than feedback inhibitory networks at decorrelating similar inputs (Wick et al., 2010). Our model incorporated both types of connectivity between SAC and EC, but we did not investigate the contributions of each type of connection; such an exploration would be an interesting direction in a future study.

“Input-tuned”, adaptive connectivities may also exist among other inhibitory networks within the OB - for example, the synaptic interactions between granule cell and mitral/tufted cells. Computational and experimental studies indicate that granule cell structural and synaptic plasticity supports the ability of the granule cell – mitral/tufted cell network to maintain output discriminability across changing input sets and following associative learning (Grelat et al., 2018; Li et al., 2018; Sailor et al., 2016). Furthermore, integration of new inhibitory connections through granule cell neurogenesis serves to refine odor representations (Huang et al., 2016; Lepousez et al., 2013; Li et al., 2018; Moreno et al., 2009). While these two layers of the OB network may affect outputs similarly, differences in their connectivity could contribute to the ability of the network to adapt to changing input environments on different timescales or to meet different behavioral requirements.

How such selective connectivity could arise remains an open question. Activity-dependent, inhibitory interglomerular connectivity could be accomplished in the mammalian OB through multiple mechanisms. Anatomical connections could be refined in an activity-dependent fashion, as they are for intrabulbar tufted cell projections (Marks et al., 2006). The dopaminergic phenotype and physiology of SACs are also modulated in an activity-dependent manner (Saino-saito et al., 2004; Chand et al., 2015); such plasticity could result in a redistribution of inhibitory connection strength that produces preferential inhibitory connectivity depending on receptive field tuning, as has been found in the visual cortex (Znamenskiy et al., 2018). Integrating learning rules into the SAC network model - as well as incorporating additional circuit elements such as feedforward, intraglomerular inhibition (Carey et al., 2015) - would be useful in exploring these possibilities in order to guide further experiments aimed at understanding the logic of inhibitory circuits in the OB circuits and their role in odorant coding.

## Acknowledgements

This work was supported by the National Science Foundation (DMS-1148230 and DMS-1853673) and the National Institutes of Health (1R01NS109979-01, R01DC06441 and F32DC016536). We thank Adam Puche and Michael Shipley for their helpful comments on the manuscript. The support and resources from the Center for High Performance Computing at the University of Utah are gratefully acknowledged.

## Notes

Conflict of interest: The authors declare no competing financial interests.

### Competing Interest Statement

The authors have declared no competing interest.

